# Development of a non-invasive diagnostic method for pathogenic RNA viruses using sebum wiped from the cat’s body surface

**DOI:** 10.1101/2025.10.15.682697

**Authors:** Yuri V Fukushima, Namiko Saito, Hirohisa Mekata, Akatsuki Saito

## Abstract

The development of non-invasive diagnostic methods for zoonotic viral infections is increasingly important for both animal welfare and public health. Sebum-based diagnostic methods using commercial oil-blotting films have been applied to SARS-CoV-2 detection in humans, yet equivalent strategies for veterinary use remain largely unexplored. Severe fever with thrombocytopenia syndrome (SFTS), caused by the SFTS virus (SFTSV), presents a major health threat in Asia—especially in Japan, where multiple cases of cat-to-veterinarian transmission have been reported. To address the need for safer diagnostics, we sought to establish a sebum-based RNA virus detection method for cats. We designed primers that efficiently detected RNA from feline sebum while distinguishing it from human and feline DNA/RNA. Using this platform, we determined the ear to be the optimal sebum collection site and confirmed that feline immunodeficiency virus (FIV) RNA can be reliably identified from ear sebum with sensitivity comparable to conventional blood-based testing. In addition, we detected SFTSV RNA from sebum samples of infected cat. Our findings introduce a minimally invasive, safe diagnostic platform for feline viral infections, reducing animal distress while safeguarding veterinarians and pet owners from zoonotic risks. This strategy marks an important step toward realizing the One Health framework by advancing the well-being of both animals and humans.

## Introduction

Zoonotic diseases remain a pressing global health challenge, with numerous ribonucleic acid (RNA) virus infections—including severe acute respiratory syndrome coronavirus 2 (SARS-CoV-2), severe fever with thrombocytopenia syndrome (SFTS), Nipah virus infection, and rabies—continuing to pose serious worldwide threats [1–4]. Prompt and accurate diagnosis is essential for controlling these infections. Currently, the standard diagnostic approach relies on detecting viral RNA from blood samples. However, this method places a considerable burden on patients due to the pain and anxiety of venipuncture, as well as the risk of bleeding in individuals with hemorrhagic disorders. In addition, handling blood samples exposes healthcare personnel to hazards contracting the target virus and other blood-borne pathogens, including human immunodeficiency virus (HIV) [5, 6] and hepatitis B virus (HBV) [7], which represent serious occupational concerns.

Although less invasive diagnostic methods, such as saliva-based testing for SARS-CoV-2 infection, have been introduced [8], their use is still confined to a limited set of infectious diseases. Consequently, there is an urgent need for novel approaches that are both less invasive for patients and safer for healthcare providers. In response, efforts have been made to develop blood-free testing methods for RNA virus detection. A recent study successfully demonstrated the presence of SARS-CoV-2 RNA in facial sebum using RNA monitoring technology [9]. This method collects sebum with commercially available oil-blotting films and analyzes the extracted mRNA. Sebum-based diagnostics thus represent a promising and minimally invasive strategy, with considerable potential for refinement and broader application.

SFTS, caused by the SFTS virus (SFTSV), is a tick-borne zoonotic disease with a high fatality rate. It is currently endemic in several Asian countries, including Japan, China, South Korea, and Thailand [10]. While tick bites represent the main route of infection [11], increasing evidence has revealed that both human-to-human [12–14] and animal-to-human [15–17] transmissions exist as the infection route. SFTSV has been detected at high concentrations in various body fluids, including blood, saliva, and urine [18].

Cats, as close companion animals to humans, require protection from viral infections for both feline and public health. Infected cats also pose transmission risks to veterinarians and animal healthcare workers. Several studies have shown that cats are highly susceptible to SFTSV [18], raising concerns not only about cat-to-human transmission but also about occupational exposure during routine veterinary practices such as blood collection. Indeed, fatal cases of cat-to-human SFTSV transmission have been reported in Japan [19]. Therefore, developing safer diagnostic methods for cats is of critical importance. Building on sebum-based detection methods established in humans, we hypothesized that a diagnostic method with oil-blotting films could provide a less invasive option for cats while also reducing infection risks for veterinarians and animal nurses.

In addition to SFTSV, we also examined feline immunodeficiency virus (FIV), another major RNA virus affecting cats globally [reviewed in 20]. FIV is one of the most common and consequential feline infectious diseases, particularly in outdoor populations [21]. Transmission occurs primarily through bite wounds from infected cats [22]. Once infected, FIV targets the immune system, leaving cats susceptible to secondary infections [reviewed in 23]. Although many FIV-positive cats may remain asymptomatic for years [24, 25], immune suppression heightens vulnerability to opportunistic infections caused by otherwise harmless microbes [reviewed in 23]. FIV presents particular challenges in animal shelters, where rescued cats are often housed together, creating a higher risk of transmission [26]. Beyond medical concerns, FIV also affects adoption prospects, as households with uninfected cats may hesitate to adopt FIV-positive cats. Given these issues, early and efficient detection is essential for prevention, and the sebum-based diagnostic method proposed in this study holds substantial promise in addressing these needs.

In this study, we aimed to establish a diagnostic method for detecting feline RNA virus infections using oil-blotting films. We successfully detected FIV RNA in sebum samples and demonstrated that the sensitivity of this sebum-based method was comparable to standard blood-based testing. In addition, we detected SFTSV RNA using sebum collected with oil-blotting films. These results suggest that sebum sampling offers a promising, minimally invasive approach for diagnosing viral infections in cats, improving veterinary safety and lowering the risk of cat-to-human transmission.

## Results

### Design and evaluation of primers for the specific detection of cat housekeeping gene mRNA from body surface lipids (BSLs)

To confirm that the oil-blotting films collected sufficient BSLs, we designed primers targeting cat housekeeping gene mRNA with high sensitivity and specificity. Because films might be contaminated by examiners handling them without gloves, we also sought to develop primers capable of distinguishing cat RNA from human DNA or RNA. Genomic DNA and total RNA were extracted from human- and cat-derived cell lines, and primers for six housekeeping genes were evaluated by quantitative real-time reverse transcription polymerase chain reaction (qRT-PCR). Among these, primers targeting *Sdha* and *B2M* consistently produced threshold cycle (Ct) values at least eight cycles lower for cat RNA than for human DNA/RNA or cat genomic DNA **(Fig. 1A)**. Both *Sdha* and *B2M* also showed relatively low overall Ct values, making them suitable for practical use **(Supplementary Table 1)**. Based on these results, *Sdha* and *B2M* were selected for further testing.

**Figure 1.**
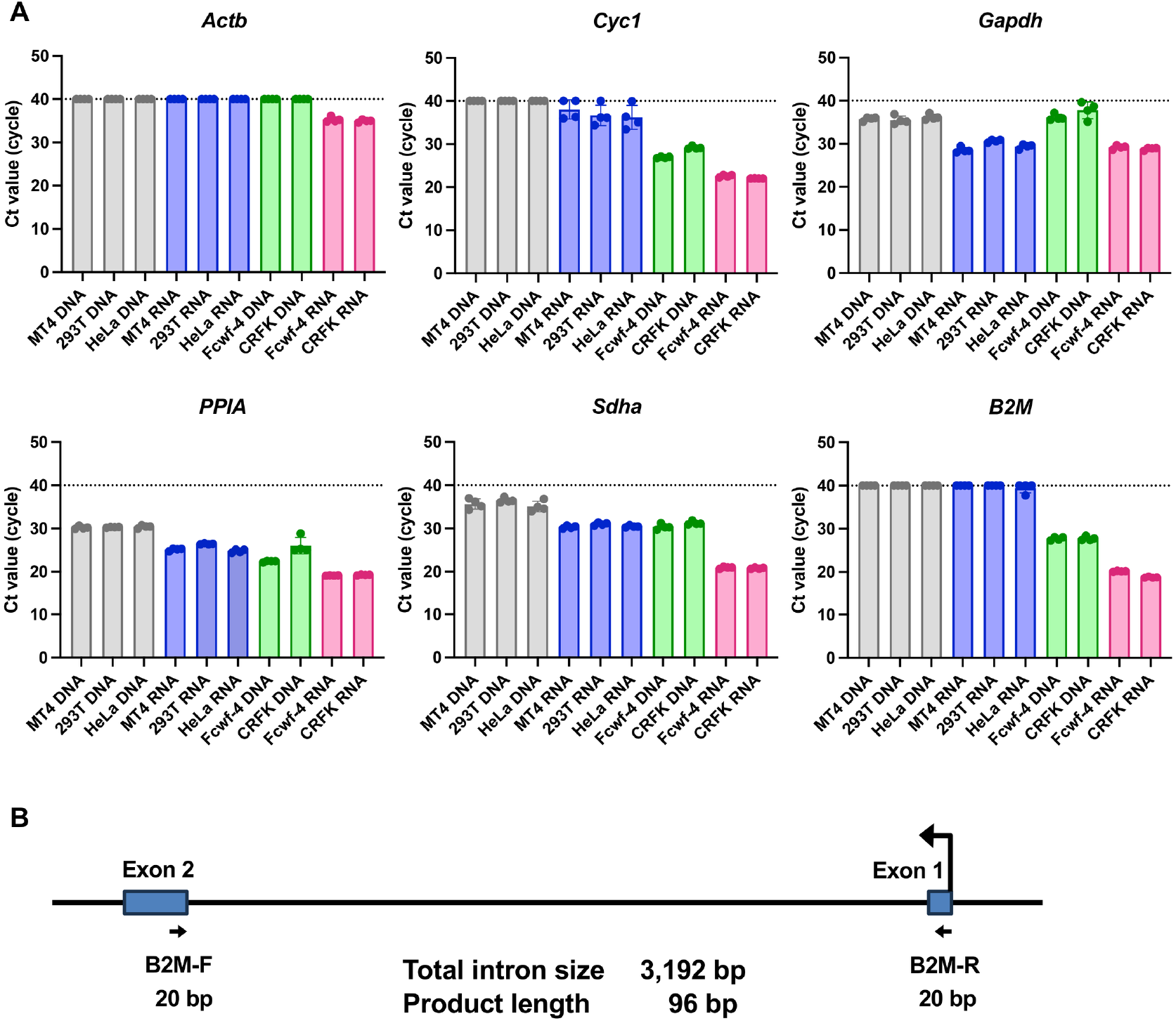
Evaluation of primers for selective amplification of the cat housekeeping gene mRNA. (A) Comparison of Ct values obtained from human genomic DNA (gray), human RNA (blue), cat genomic DNA (green), and cat RNA (red). Data show the mean ± standard deviation (SD) of quadruplicate measurements and represent at least four independent experiments. (B) Schematic of primer design. Blue boxes represent exons, arrows indicate primer sites, and F and R mark forward and reverse primers, respectively.

We next examined whether transcripts of *Sdha* and *B2M* could be detected from BSLs using the designed primers **(Fig. 1A)**. Four oil-blotting film samples obtained from three cats were analyzed. *B2M* was successfully detected with an average Ct value of 36.2 **(Fig. 2A)**, whereas *Sdha* was not detectable from BSLs **(Fig. 2B)**. On this basis, *B2M* was chosen as the optimal primer, and a probe was subsequently designed for *B2M* to enable probe-based detection **(Fig. 1B, Supplementary Table 2)**.

**Figure 2.**
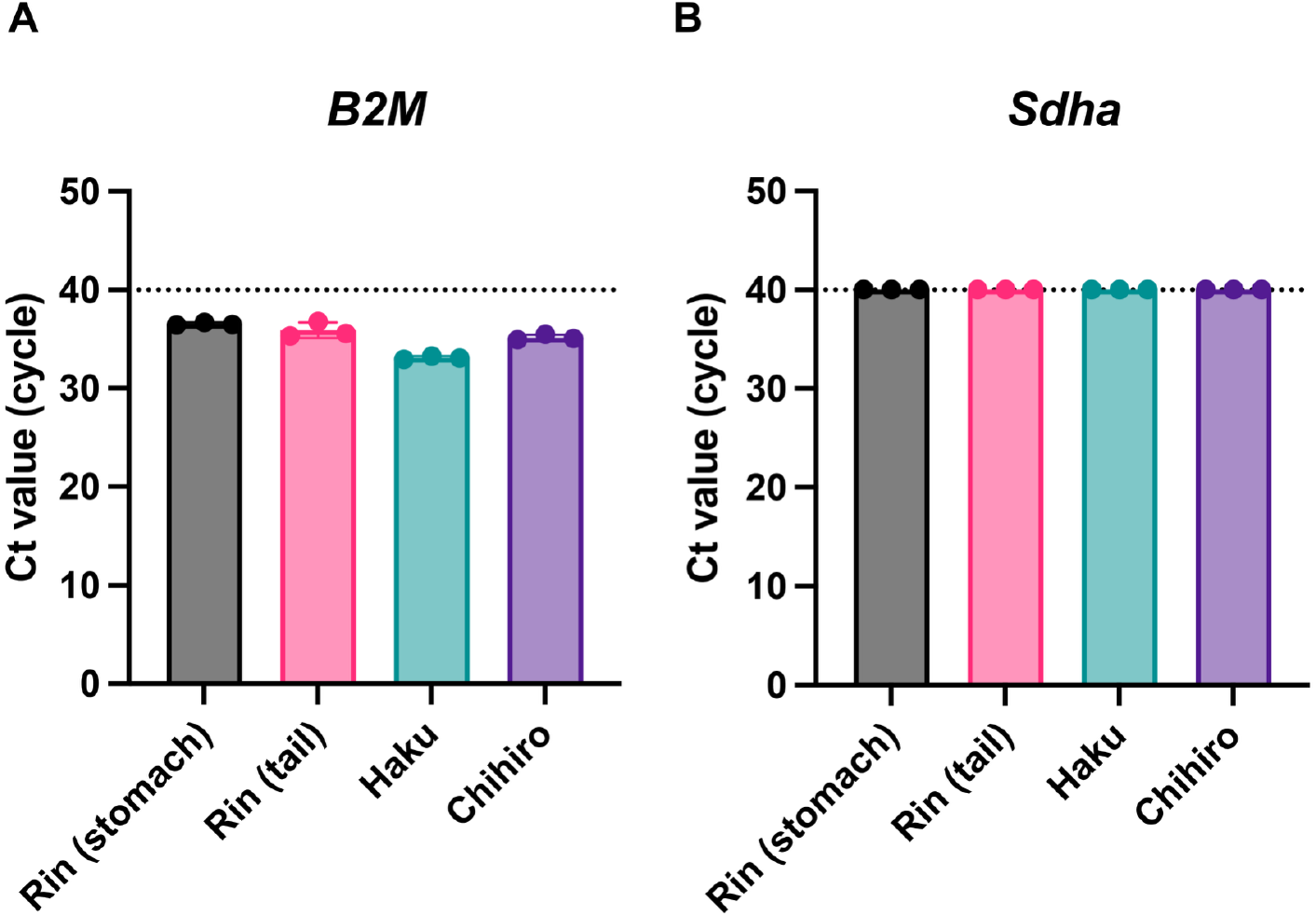
Detection of cat housekeeping gene mRNA from body surface lipids (BSLs). Four BSL samples were collected from three cats. Data show the mean ± SD of triplicate measurements from one assay, representative of at least three independent experiments.

### Optimization of sample processing for sensitive RNA detection from cat BSLs

We first determined the optimal sampling site for RNA collection from cat BSLs. Samples were collected from the armpit, tail base, tail, and ear of the same cat. The tail base, tail, and ear all yielded significantly lower Ct values for *B2M* compared to the armpit (**Fig. 3A**). Because the subject was an intact adult male, known to secrete higher levels of BSLs [27], we concluded that the ear represents the most reliable sampling site across sexes and age groups. All subsequent experiments were therefore conducted using ear-derived BSLs (**Fig. 3B–D**).

**Figure 3.**
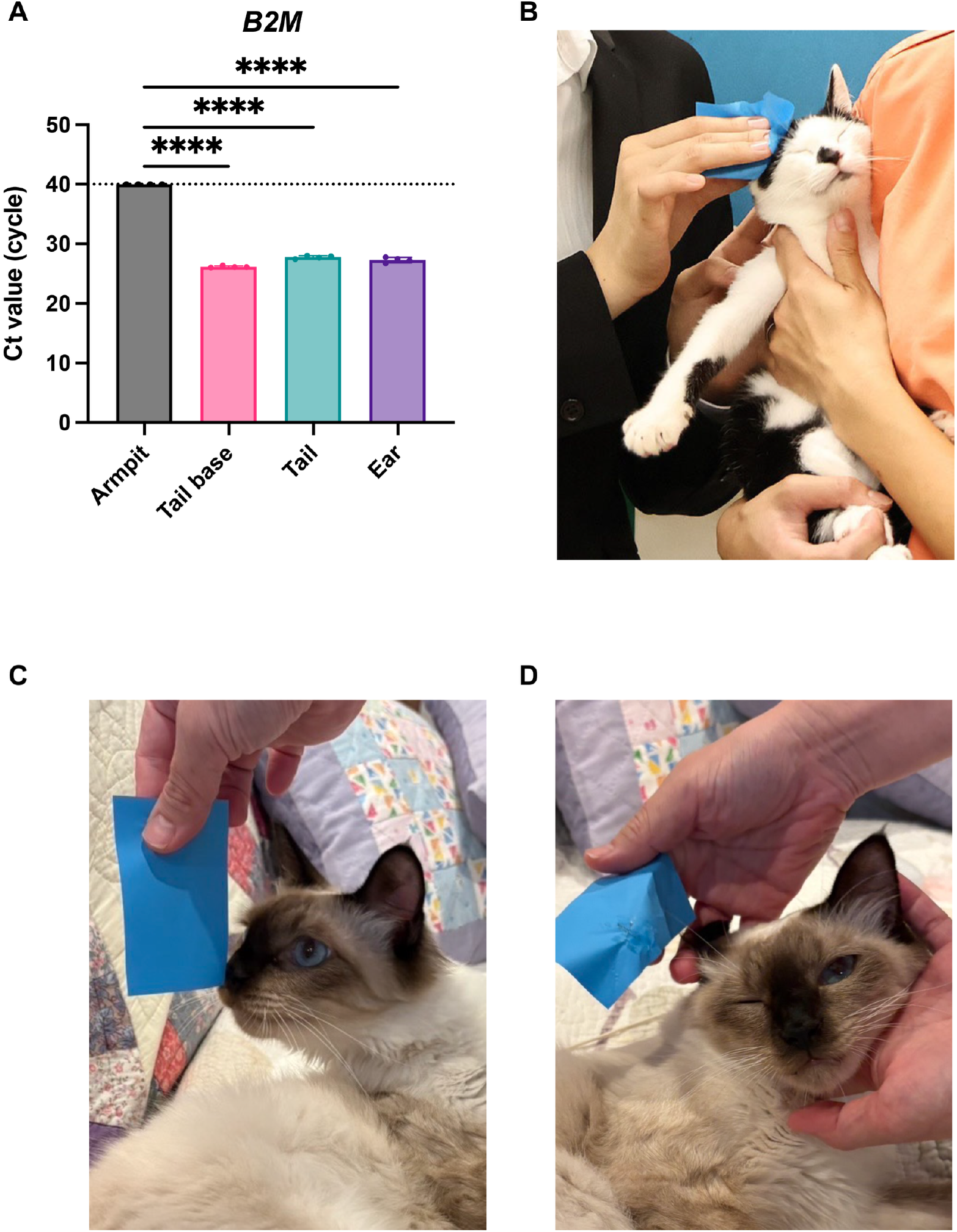
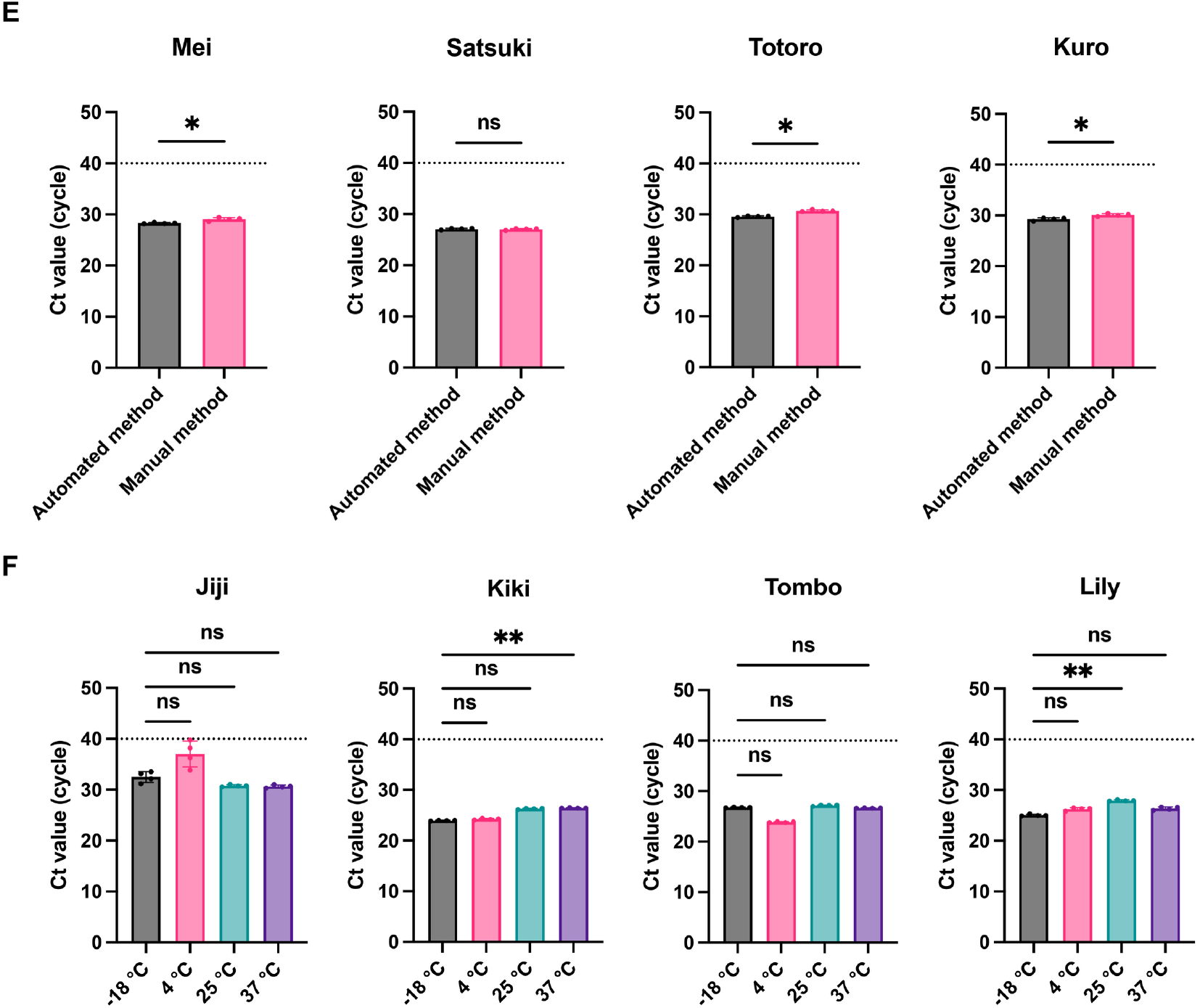
Optimization of sample handling for sensitive RNA detection from cat body surface lipids (BSLs). (A) Ct values of *B2M* measured in BSLs from four sampling sites in the same cat. Data show the mean ± SD of quadruplicate measurements. Differences were analyzed by one-way ANOVA using armpit-derived BSLs as the control; \*\*\**p* < 0.001, \*\**p* < 0.01, \**p* < 0.05, ns = not significant. Representative image of sebum collection from a cat’s side. (C) Oil-blotting film before sampling. (D) Oil-blotting film after sampling, with darker areas indicating collected sebum. (E) Comparison of *B2M* detection sensitivity between manual and automated RNA extraction. Data show the mean ± SD of quadruplicate measurements. Differences were analyzed by one-way ANOVA using armpit-derived BSLs as the control; \*\*\**p* < 0.001, \*\**p* < 0.01, \**p* < 0.05, and ns = not significant. (F) Effect of storage temperature on *B2M* detection in BSLs stored on oil-blotting films at −18 °C, 4 °C, 25 °C, or 37 °C for three days. Data show the mean ± SD of quadruplicate measurements, representative of at least four experiments. Statistical significance was determined by one-way ANOVA; \*\*\**p* < 0.001, \*\**p* < 0.01, \**p* < 0.05, ns = not significant.

For practical application, we further evaluated whether RNA extraction could be fully automated. Samples from four cats were processed using either a manual protocol (QIAzol + Viral RNA Mini Kit) or a fully automated extraction system (All Maxwell). The automated method provided higher B2M detection sensitivity than the manual protocol (**Fig. 3E**). Owing to its simplicity and high sensitivity, we adopted the automated method for subsequent experiments. Finally, we tested the stability of BSL samples under different storage conditions to simulate clinical and transport scenarios. Oil-blotting films containing BSLs were stored at −18 °C, 4 °C, 25 °C, and 37 °C for three days, reflecting estimated maximum shipping times in Japan. Samples from four cats showed no significant differences in *B2M* detection across the tested temperatures (**Fig. 3F**).

### Efficient detection of FIV virus RNA from BSLs

After optimizing the sampling site, we evaluated the feasibility of detecting B2M and FIV RNA from ear-derived BSLs. Samples were collected from 12 cats. *B2M* was successfully detected in all samples (**Fig. 4A**), confirming the ear as the optimal site for BSL collection. FIV RNA was detected only in cats previously diagnosed as FIV-positive by blood testing (**Fig. 4B**), demonstrating full agreement between sebum-based and conventional diagnostics.

**Figure 4.**
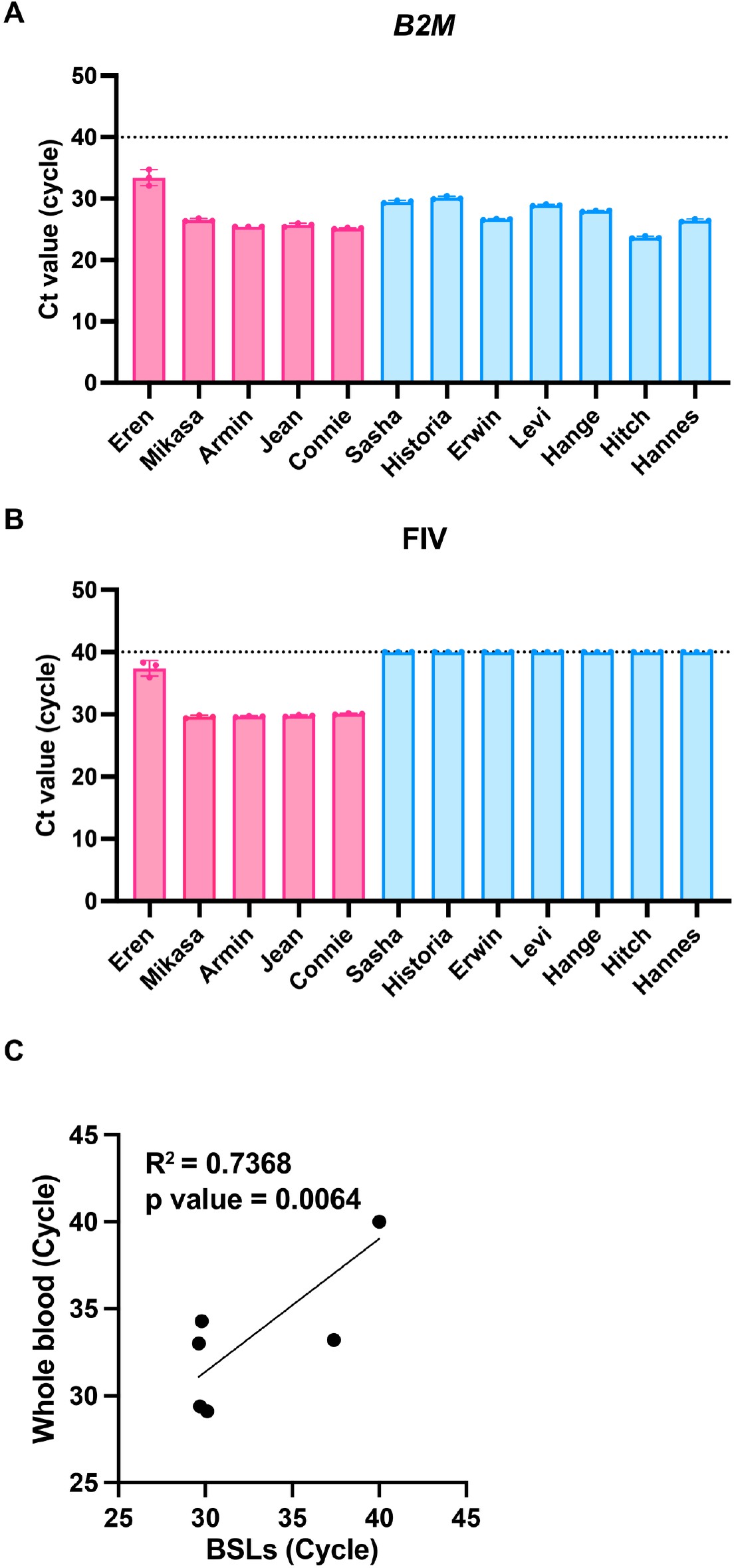
Efficient detection of feline immunodeficiency virus (FIV) RNA from body surface lipids (BSLs). (A) Ct values of *B2M* from BSLs collected from 12 cats. Red symbols indicate FIV-positive cats; blue symbols indicate FIV-negative cats. Data show the mean ± SD of triplicate measurements. (B) Ct values of FIV RNA detected in the same BSL samples. (C) Correlation of Ct values obtained by sebum-based versus conventional blood-based testing for FIV RNA.

Using values shown in **Fig. 4B**, we compared the performance of the sebum-based diagnostic method with the blood-based method in terms of FIV RNA sensitivity. Oil-blotting film and whole-blood samples were collected from the same six cats, and Ct values for FIV RNA were analyzed. The oil-blotting film method showed detection sensitivity equivalent to that of conventional whole-blood testing (**Fig. 4C**).

### Detection of SFTSV RNA from cat BSLs

As the final experiment, we investigated whether SFTSV RNA could be detected from sebum. BSLs were collected with oil-blotting films from the base of the tail of a cat naturally infected with SFTS. SFTSV RNA was successfully detected only from BSL samples collected at the base of the tail (**Fig. 5**).

**Figure 5.**
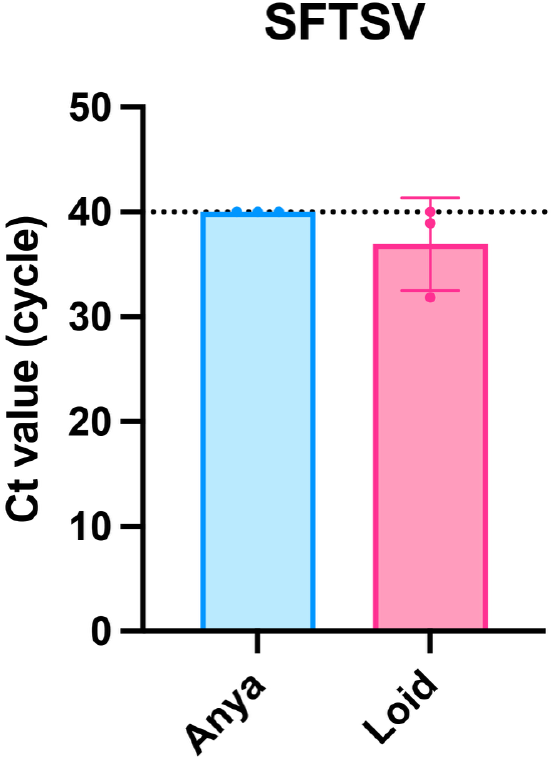
Detection of severe fever with thrombocytopenia syndrome virus (SFTSV) RNA from cat body surface lipids (BSLs). Ct values of SFTSV measured in BSLs from two cats. Red symbols represent the SFTSV-positive cat (Loid), and blue symbols represent the SFTSV-negative cat (Anya).

## Discussion

In this study, we established a sebum-based diagnostic method for detecting RNA virus infections in cats. This approach aimed to minimize invasiveness for animals while enhancing safety for examiners. We developed primers that efficiently amplified viral RNA from feline sebum while distinguishing it from human DNA/RNA and feline genomic DNA. We also identified optimal sampling sites and showed that FIV RNA, a major feline pathogen, could be reliably detected from ear-derived sebum. Notably, the sensitivity of FIV detection using sebum was equivalent to that of conventional blood-based testing. Moreover, we successfully detected SFTSV RNA from sebum collected at the tail base, providing proof of concept for sebum-based diagnostics of SFTS.

The establishment of a sebum-based diagnostic method has important implications for both animal welfare and the One Health framework. By substituting needle-based blood collection with non-invasive sebum sampling, the physical and psychological burden on cats can be substantially reduced. This method aligns with the Five Freedoms of animal welfare, particularly “Freedom from Pain, Injury, or Disease” and “Freedom from Fear and Distress.” In addition, sebum-based testing could improve biosafety for cat owners, local communities, veterinarians, and veterinary nurses by lowering the risk of exposure to infectious agents during sample handling.

Previous studies have demonstrated that sebum contains diverse biomolecules, including mRNA, as a result of holocrine secretion in human [28]. On this basis, we hypothesized that feline sebum would also harbor mRNA. Supporting this, we successfully detected *B2M*-specific mRNA from feline sebum (**Fig. 2**), establishing a biological rationale for RNA detection from this sample type. To determine optimal sampling sites, we compared *B2M* Ct values across multiple body regions and found that ear-derived sebum consistently provided one of the strongest detection signals (**Fig. 3A, 3B, 3C, 3D**). Considering potential variation by sex and age, we focused subsequent experiments on ear sampling, which yielded reproducible RNA detection outcomes.

To enable practical application, we incorporated automated RNA extraction (**Fig. 3E**), which markedly shortened processing time without compromising sensitivity. We further evaluated sample stability across diverse storage conditions. *B2M* detection sensitivity remained consistent after three days at temperatures ranging from −18 °C to 37 °C (**Fig. 3F**). These results support the feasibility of nationwide sample shipment, potentially expanding diagnostic access and mitigating healthcare disparities, particularly in geographically remote regions.

Regarding FIV diagnostics, *B2M* mRNA was identified in all samples, ensuring ear-derived sebum as a reliable source for sebum-based testing (**Fig. 4A**). Sebum- and blood-based diagnostics results showed complete concordance (**Fig. 4B**). Moreover, FIV detection with sebum-based method achieved sensitivity equivalent to conventional blood testing (**Fig. 4C**), underscoring its promise for clinical adoption. Unlike antibody-based kits, which cannot distinguish maternal antibodies from true infection [29], this method directly targets viral RNA, enabling accurate diagnosis in kittens. Collectively, these findings indicate that sebum-based diagnostics could augment or strengthen current testing strategies.

Importantly, this work presents the first evidence that SFTSV RNA is detectable in feline sebum (**Fig. 5**), providing proof-of-concept for sebum-based SFTS diagnostics. Subsequent studies will compare detection sensitivity between sebum and blood samples and assess clinical utility across broader populations.

Despite encouraging outcomes, blood testing remains indispensable for a comprehensive health assessment. Hematological measures, such as platelet and white blood cell counts in SFTS [30] and CD4 T-cell counts in FIV [31], are critical for diagnosis. We therefore propose an integrated framework in which sebum- and blood-based tests are applied complementarily, depending on clinical objectives. Sebum testing is especially useful when venipuncture is difficult—for example, in fearful or aggressive cats, large-scale screening of community cats, or suspected zoonotic risk cases. It also provides a safer option for kittens and clinically fragile animals.

For cats prone to severe stress during veterinary visits, blood collection can worsen anxiety and deter future care [reviewed in 32]. By contrast, sebum collection from the ear requires only gentle restraint and produces substantially less stress. Owners could also be trained to collect sebum at home, enhancing access to diagnostic testing for cats that are hard to manage in clinics.

In this study, numerous samples were collected from cats housed at the Miyazaki City shelter. Owing to its simplicity, safety, and low training threshold, sebum-based testing is particularly suited to high-throughput shelter screening. Unlike blood collection, which generally requires two trained professionals, ear-derived sebum sampling can be performed by one individual with limited instruction. Considering the global importance of FIV and FeLV screening in shelters [26], this method offers considerable value for preventing viral spread before adoption. Ongoing validation studies combining sebum- and blood-based diagnostics are underway in collaboration with Miyazaki City shelters, aiming to establish standardized protocols.

This study has several limitations. First, sebum-based testing focuses on pathogen detection and cannot substitute for blood tests in evaluating hematological or biochemical markers. Second, in multi-cat environments, behaviors such as ear-licking may introduce saliva contamination, potentially yielding false positives, especially for viruses present in body fluids such as SFTSV [18]. This limitation may be alleviated by gathering behavioral information from owners and integrating targeted screening questions. Finally, the relatively complex RNA extraction workflow remains a practical barrier. To address this, we developed a fully automated extraction platform (**Fig. 3E**), facilitating wider clinical and epidemiological application.

In conclusion, the sebum-based diagnostic method developed in this study offers a minimally invasive and highly reliable approach for detecting RNA viral infections in cats. Applying this technique in practice may also enhance examiner safety by lowering risks linked to blood handling and reducing direct contact with cats potentially carrying zoonotic pathogens. Together, these advantages emphasize the broad utility of sebum-based diagnostics for clinical practice, shelter management, and epidemiological surveillance, thereby supporting the One Health framework. Future work will focus on refining sampling protocols, validating diagnostic performance across diverse populations, and establishing standardized guidelines for veterinary and public health professionals. Collectively, these efforts aim to strengthen feline viral infection control, improve animal welfare, and promote wider adoption of minimally invasive and safe diagnostic tools for RNA virus detection.

## Methods

### Ethical approval

All procedures involving the collection of feline samples were approved by the Animal Care and Use Committee of the University of Miyazaki (approval No. 2025-024) and conducted in accordance with the University of Miyazaki Experimentation regulations and relevant institutional guidelines.

### BSL sample collection

BSLs were collected from animal hospitals and the Miyazaki Animal Protection Center between March and August 2025 using STF Oil Clear Film (Hakugen Earth Co., Ltd., Tokyo, Japan; 75 mm × 160 mm × 3 mm; JAN code: 4902407040930). Sampling was performed by gentle wiping designated areas, without anesthesia or physical restraint. Each used film was sealed individually in a plastic bag and stored at −30 °C until processing.

### Cell culture

Lenti-X 293T cells (*Homo sapiens*; TaKaRa Bio Inc., Kusatsu, Japan; Cat# Z2180N), HeLa cells (*Homo sapiens*; ATCC, Manassas, VA, USA; Cat# CCL-2), Fcwf-4 cells (*Felis catus*; ATCC; Cat# CRL-2787), and Crandell-Rees feline kidney (CRFK) cells (*Felis catus*; JCRB, Ibaraki, Japan; Cat# JCRB9035) were maintained in Dulbecco’s modified Eagle’s medium (Nacalai Tesque, Kyoto, Japan; Cat# 08458-16) supplemented with 10% fetal bovine serum (FBS) and 1× penicillin– streptomycin (Nacalai Tesque; Cat# 09367-34). MT4 cells (*Homo sapiens*; JCRB; Cat# JCRB0135) were maintained in Roswell Park Memorial Institute medium (RPMI-1640; Nacalai Tesque; Cat# 30264-56) with 10% FBS and 1× penicillin–streptomycin (Nacalai Tesque; Cat# 09367-34).

### Extraction of RNA from cat BSLs

Sebum samples collected on oil-blotting films were cut into small pieces and placed in 2.0 mL microcentrifuge tubes. QIAzol reagent (1,450 µL; QIAGEN, Hilden, Germany; Cat# 79306) was added, and tubes with samples were incubated in a constant-temperature shaker at 200 rpm for 5 min at room temperature. After brief vortexing, 1,300 µL of the supernatant was transferred to fresh tubes. Chloroform (260 µL; Nacalai Tesque, Kyoto, Japan; Cat# 08402-84) was added, followed by vortexing and centrifugation at 15,000 × g for 15 min at 4°C. The aqueous phase (600 µL) was carefully collected, and RNA was purified with the RNeasy Mini QIAcube Kit (QIAGEN; Cat# 74116), including on-column DNase treatment under a custom protocol. In brief, 600 µL of 85% ethanol was added, which were mixed thoroughly and loaded onto RNeasy Mini spin columns. The columns were washed with RW1 buffer, treated with DNase I, washed again with RW1, and finally washed with RPE buffer. RNA was eluted twice with 50 µL RNase-free water, yielding a final volume of 100 µL.

RNA cleanup was performed using the Monarch® Spin RNA Cleanup Kit (10 µg; New England Biolabs, Ipswich, MA, USA; Cat# T2030S). In brief, 200 µL Buffer BX and 300 µL ethanol were added to 100 µL RNA solution and mixed by pipetting. The mixture was loaded onto the spin column and centrifuged at maximum speed for 1 min. The column was then washed twice with 500 µL Buffer WX, each step followed by centrifugation for 1 min. After a final dry spin to remove residual buffer, the column was placed onto a new RNase-free 1.5 mL microcentrifuge tube, and RNA was eluted with 8 µL RNase-free water.

For automated RNA extraction, RNA from cat BSLs was processed using the Maxwell® RSC 48 Instrument (Promega, Madison, WI, USA; Cat# AS8500) in combination with the Maxwell® RSC miRNA Plasma and Serum Kit (Promega; Cat# AS1680), according to the manufacturer’s instructions.

### Primers

Primers were designed with reference to earlier studies [33] and the QIAGEN housekeeping gene database [34]. Sequences were generated using Primer-BLAST (https://www.ncbi.nlm.nih.gov/tools/primer-blast/; accessed May 5, 2025) based on the *Felis catus* genomic sequence (accession no. JAFEKA010000007.1). The following criteria were applied for ideal primer design: (1) expected PCR product size was set between 70–150 bp, (2) primers separated by at least 1,000 bp of intronic sequence, and (3) primers were required to span an exon–exon junction **(Supplementary Table 2)**. All primers were synthesized as standard desalted DNA oligonucleotides by Eurofins Genomics (Tokyo, Japan) and stored at 4 °C until use.

### Real-time quantitative polymerase chain reaction (RT-qPCR)

Total DNA was extracted from cultured cells using the DNeasy Blood & Tissue Kit (QIAGEN; Cat# 69506) with RNase A (17,500 U; QIAGEN; Cat# 19101) and 100% ethanol, following the manufacturer’s protocol. Total RNA was extracted from cultured cells using the RNeasy Mini Kit (QIAGEN; Cat# 74104) with QIAshredder columns (QIAGEN; Cat# 79656), the RNase-Free DNase Set (QIAGEN; Cat# 79254), and 2-mercaptoethanol (Bio-Rad, Hercules, CA, USA; Cat# 1610710). RNA from sebum samples on oil-blotting films was purified with the Monarch® Spin RNA Cleanup Kit with 85%–100% ethanol. DNA and RNA concentrations were measured using a NanoDrop 2000 spectrophotometer (Thermo Fisher Scientific, Waltham, MA, USA).

The mRNA expression of six housekeeping genes—*Actin Beta (Actb), Cytochrome C1 (Cyc1), glyceraldehyde-3-phosphate dehydrogenase (Gapdh), peptidylprolyl isomerase A (PPIA), succinate dehydrogenase complex flavoprotein subunit A (Sdha)*, and *Beta-2-microglobulin (B2M)*—was quantified using the One Step TB Green® PrimeScript™ PLUS RT-PCR Kit (Perfect Real Time) (TaKaRa Bio, Shiga, Japan; Cat# RR096A). RT-qPCR was carried out under the following cycling conditions: reverse transcription at 42 °C for 5 min, initial denaturation at 95 °C for 10 s, followed by 40 cycles of 95 °C for 5 s and 60 °C for 34 s.

To quantify FIV RNA (**Supplementary Table 3**), SFTSV RNA (**Supplementary Table 4**), and *B2M* mRNA, RT-qPCR was performed using the PrimeDirect® Probe RT-qPCR Mix (TaKaRa; Cat# RR650A). Conditions were reverse transcription at 90 °C for 3 min, annealing at 60 °C for 5 min, and 40 cycles of 95 °C for 5 s and 60 °C for 30 s. For all reactions, melting-curve analysis was conducted to verify amplification specificity, and relative expression levels were determined using the 2^^ΔCt^ method.

### Statistical analysis

The differences in Ct values across treatments were assessed using one-way analysis of variance (ANOVA), followed by Tukey’s post hoc test, or by the Kruskal–Wallis test with Dunn’s post hoc test where appropriate. A *p*-value ≤ 0.05 was considered statistically significant. All analyses were conducted in GraphPad Prism (ver. 10.2.3; GraphPad Software, San Diego, CA, USA).

## Acknowledgments

The authors thank Ms. Saori Kusakari, Ms. Tomoko Nishiuchi, Ms. Miki Kawano, Ms. Natsumi Matsubara, Ms. Kana Shimada, Ms. Yuka Oshikawa, Ms. Yukino Tokito, Ms. Kaoru Terashi, clinical veterinarians, the staff of CADIC, University of Miyazaki, the staff of Miyazaki Animal Welfare Center (Miyazaki City), and all the cats involved in this study, especially Nanako, Robin, Chacha, Aka, Maguro, Kahlúa, Ando, and Ruru-ta, for their assistance.

## Author information

### Contributions

YV.F., H.M., and A.S. designed the experiments. YV.F., N.S., H.M., and A.S. performed the experiments. YV.F., H.M., and A.S. analyzed the results. YV.F., H.M., and A.S. wrote the manuscript. All authors have read and approved the manuscript.

### Data availability

Source data are available on request.

