## Supplemental Information for "Development of a non-invasive diagnostic method for pathogenic RNA viruses using sebum wiped from the cat’s body surface"

**Supplementary Table 1. Ct value average of Fcwf-4 RNA and CRFK RNA**

| | ***Actb*** | ***Cyc1*** | ***Gapdh*** | ***PPIA*** | ***Sdha*** | ***B2M*** |
|---|---|---|---|---|---|---|
| **Ct value** | 35.2 | 22.3 | 29.0 | 19.2 | 20.9 | 19.4 |

**Supplementary Table 2. Sequences of primers/probe to amplify cat housekeeping genes**

| Target gene | | Sequence (5'->3') | Tm | Length | Amplicon size |
|---|---|---|---|---|---|
| *Actb* | **Forward** | CGGCGCCGCCCTATAAA | 59.9 | 17 | 144 |
| | **Reverse** | TCATCATCCATGGCGAACTGAG | 60.5 | 22 | |
| *Cyc1* | **Forward** | TTGGAGTATGACGATGGCACC | 60.1 | 21 | 149 |
| | **Reverse** | AGAGGCAATAGCAAGCCCAT | 59.5 | 20 | |
| *Gapdh* | **Forward** | TGAAGGTCGGTGTGAACGG | 59.9 | 19 | 134 |
| | **Reverse** | CTGGAACATGTAGACCATGTAGT | 57.7 | 23 | |
| *PPIA* | **Forward** | GGTCAACCCCATCGTGTTTTT | 59.3 | 21 | 79 |
| | **Reverse** | GTCTGCAAACAGGTCGAAGG | 59.1 | 20 | |
| *Sdha* | **Forward** | CACTGACTAGGGCGCAGTGG | 62.8 | 20 | 140 |
| | **Reverse** | AAATTCATGGTCCACTACCGGG | 60.4 | 22 | |
| *B2M* | **Forward** | GCGTTTTGTGGTCTTGGTCC | 63.7 | 20 | 96 |
| | **Reverse** | GGGTGACGGGAGTAAACCTG | 63.9 | 20 | |
| | **Probe** | /5HEX/TGGATGCCG/ZEN/TCCAGCATTCTCCA/3IABkFQ/ | 69.7 | 23 | |

**Supplementary Table 3. Sequences of primers and probe to amplify FIV**

| | Sequence (5'->3') | Tm | Length | Amplicon size |
|---|---|---|---|---|
| **Forward** | GCCTTCTCTGCAAATTTAACACCT | 63.9 | 24 | 91 |
| **Reverse** | GATCATATTCTGCTGTCAATTGCTTT | 62.7 | 26 | |
| **Probe** | /5Cy5/CATGGCCAC/TAO/ATTAATAATGGCCGCA/3IAbRQSp/ | 67.7 | 25 | |

**Supplementary Table 4. Sequences of primers and probe to amplify SFTSV**

| | Sequence (5’->3’) | Tm | Length | Amplicon size |
|---|---|---|---|---|
| **Forward** | TGTCAGAGTGGTCCAGGATT | 62.6 | 20 | 137 |
| **Reverse** | ACCTGTCTCCTTCAGCTTCT | 62.6 | 20 | |
| **Probe** | /56-FAM/TGGAGTTTG/ZEN/GTGAGCAGCAGC/3IABkFQ/ | 67.1 | 21 | |
